# Bioluminescent detection of viral surface proteins using branched multivalent protein switches

**DOI:** 10.1101/2023.09.22.558942

**Authors:** Alexander Gräwe, Cindy M. Spruit, Robert P. de Vries, Maarten Merkx

**Affiliations:** Laboratory of Protein Engineering, Department of Biomedical Engineering and Institute for Complex Molecular Systems, Eindhoven University of Technology, Eindhoven, the Netherlands; Utrecht Institute for Pharmaceutical Sciences, Department of Chemical Biology and Drug Discovery, Utrecht, the Netherlands

## Abstract

Fast and reliable virus diagnostics is key to prevent the spread of viruses in populations. A hallmark of viruses is the presence of multivalent surface proteins, a property that can be harnessed to control conformational switching in sensor proteins. Here, we introduce a new sensor platform (dark-LUX) for the detection of viral surface proteins consisting of a general bioluminescent framework that can be post-translationally functionalized with separately expressed binding domains. The platform relies on 1) plug-and-play bioconjugation of different binding proteins via SpyTag/SpyCatcher technology to create branched protein structures, 2) an optimized turn-on bioluminescent switch based on complementation of the split-luciferase NanoBiT upon target binding and 3) straightforward exploration of the protein linker space. The influenza A virus (IAV) surface proteins hemagglutinin (HA) and neuraminidase (NA) were used as relevant multivalent targets to establish proof of principle and optimize relevant parameters such as linker properties, choice of target binding domains and the optimal combination of the competing NanoBiT components SmBiT and DarkBiT. The sensor framework allows rapid conjugation and exchange of various binding domains including scFvs, nanobodies and *de novo* designed binders for a variety of targets, including the construction of a heterobivalent switch that targets the head and stem region of hemagglutinin. The modularity of the platform thus allows straightforward optimization of binding domains and scaffold properties for existing viral targets, and is well suited to quickly adapt bioluminescent sensor proteins to effectively detect newly evolving viral epitopes.

## Introduction

New strains of viruses have the potential to cause future pandemics, disrupting global healthcare and well-being as seen for SARS-CoV-2. To restrain virus spread, fast and reliable virus diagnostics are thus highly relevant. Influenza A viruses (IAV) are prime candidates for a next pandemic, due to their zoonotic nature ^1^. Current virus detection methods such as PCR rely on the identification of virus nucleic acids, which renders them sensitive but requires specialized equipment, precluding testing at the point of care (POC). Antibody-based lateral flow antigen tests are faster but do not yet reach the sensitivity of PCR ^2^. As IAVs constantly adapt, the rapid creation of new, sensitive lateral flow assays for direct viral protein detection remains challenging.

Homogeneous immunoassays based on engineered protein sensors are attractive for POC diagnostics as they enable direct analyte detection in the sample of interest ^3^. Sensor proteins that use bioluminescence as readout are of particular interest, as the absence of light scattering and background fluorescence enables measurements in complex matrices. Furthermore, detection can be done using minimal hardware, such as a standard smartphone or digital camera ^4^. Baker and co-workers recently reported a bioluminescent sensor for SARS-CoV-2 based on Bioluminescent Resonance Energy Transfer (BRET). Their FUS231-P12 sensor protein consisted of *de novo* designed binders flanked with BRET donor and acceptor domains. In this particular case, binding of the sensor to multiple adjacent domains on the SARS-CoV-2 spike protein resulted in an increase in BRET ^5^. However, the change in BRET signal depends critically on the distance between donor and acceptor in the target bound state, which will be different for each new target.

An alternative bioluminescent sensor mechanism is based on the control of the intramolecular complementation of split luciferases. This mechanism has been used in a recent version of the LUMABS family of antibody sensors that use a split version of NanoLuc called NanoBiT ^6^, consisting of a large (LgBiT) and small (SmBiT) subunit ^7–10^. In these sensors, bivalent binding to the two antigen binding domains of an antibody leads to intramolecular domain rearrangement, which results in a change of bioluminescence color (NB-LUMABS) or an increase in bioluminescence intensity (dark-LUMABS). While these sensors – like the FUS231-P12 sensor – rely on multivalent interactions, the sensor response itself does not critically depend on the exact architecture of the target, and can be optimized by tuning the relative affinities of the competing split luciferase domains.

Multivalent binding of a sensor to a target protein is beneficial as the resulting avidity effects increase the sensitivity and specificity of a sensor ^11^. Interestingly, avidity is also the key driving force that allows IAVs to bind to their target cells, compensating for the low affinities of one hemagglutinin domain (HA) to sialic acid (K_D_ = 0.5-20 mM) ^12^. The HA protein is a homotrimer, whereas the neuraminidase (NA) surface protein is present as a homotetramer. While both FUS231-P12 and LUMABS biosensors harness the multivalent nature of their targets, they have some drawbacks as dedicated sensor platforms for the detection of viral surface proteins. Firstly, they rely on peptide epitopes or *de novo* designed small binding domains, but they are less suitable to be used in combination with multiple structurally more complex binding domains such as nanobodies and scFv’s. Secondly, the designs are not created for straightforward optimization of linkers, which is often required to achieve an optimal multivalent binding between the sensor and new targets ^13,14^.

To overcome these limitations, we developed a modular platform that fulfills the requirements for the detection of multivalent viral targets by combining reliable technologies and advances in the field of protein engineering. We constructed a general sensor framework containing the NanoBiT and DarkBiT components required for switching between a ‘dark’ OFF and a blue ‘ON’ state. We further integrated the SpyCatcher bioconjugation strategy ^15^ to allow attachment of target binding domains to this scaffold, yielding fusion proteins with branches formed by isopeptide bond formation between SpyCatcher003/SpyTag003 or DogCatcher/DogTag (Fig. 1A). When the assembled sensors bind multivalent targets, spatial separation promotes dissociation of the complex between LgBiT and DarkBiT, allowing the formation of an active luciferase formed by binding of LgBiT to SmBiT and generation of a bioluminescent signal (Fig. 1B). The iFLinkC method ^16^ was used to generate a library of sensor proteins with different linker lengths and physicochemical properties and screen for linkers that achieve a higher dynamic range in signal output.

**Figure 1.**
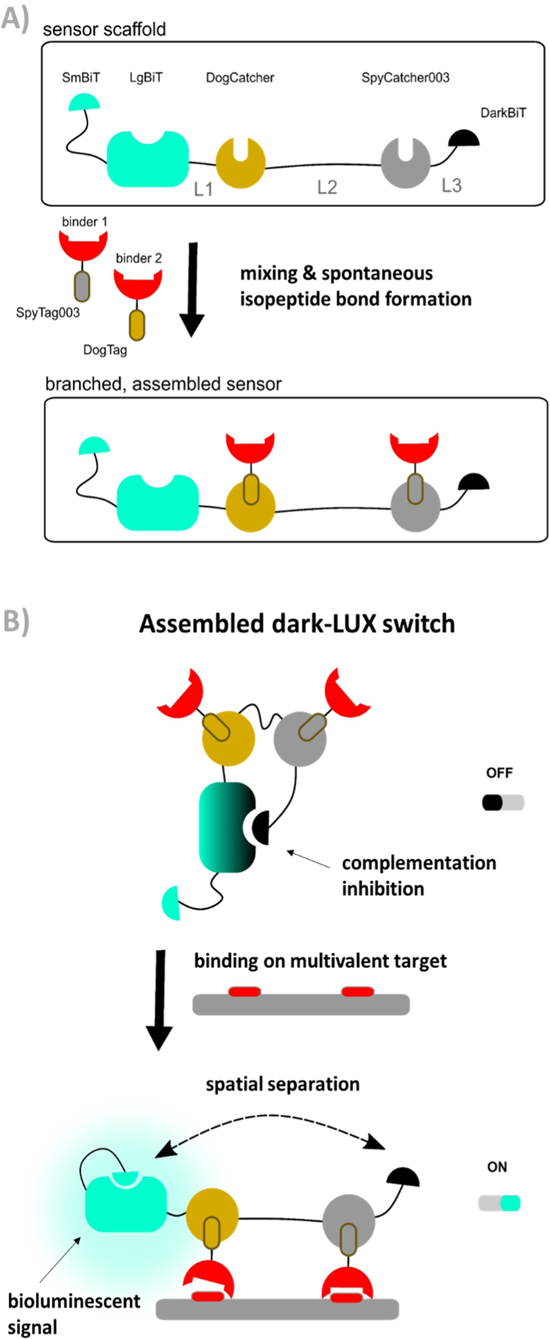
Concept of the dark-LUX sensor platform. A) Spontaneous formation of multivalent, branched dark-LUX sensors by mixing a sensor scaffold containing DogCatcher and SpyCatcher003 with separately expressed binders containing the matching DogTag and SpyTag003 tags. B) Concept of dark-LUX sensors. A fully assembled dark-LUX sensor consists of the dark-LUX scaffold and two binders. In the presence of a multivalent target, the binders bind to the target, resulting in a spatial separation of DarkBiT and LgBiT, provided the intramolecular affinities and linkers allow the conformational change. This allows SmBiT to complement LgBiT and form an active luciferase complex that generates light by conversion of furimazine.

## Results

### Design and development of the dark-LUX sensor platform

Key to the development of a sensor with good sensitivity is the suppression of background signal in the absence of target molecules. Therefore, we started by devising a sensor system that shows a minimal signal in the inactive (OFF) state. For this, we used the split-NanoLuc system NanoBiT in combination with a high-affinity DarkBiT peptide as complementation inhibitor. DarkBiTs are small (11-13 aa) peptides that differ from SmBiTs by a single R>A mutation that renders them unable to form an active NanoBiT complex but still allows binding to LgBiT^10^. To achieve switching to an ON state, we introduced a SmBiT that is able to complement LgBiT upon dissociation of the LgBiT-DarkBiT interaction. This dissociation is achieved when a conformational change increases the intramolecular distance between LgBiT and DarkBiT. We recently accomplished this for the antibody detection platform dark-LUMABS by introducing two epitopes to the sensor that are connected via a long semi-flexible linker. The linker extends when the target antibody is bivalently bound, providing the necessary spatial separation of LgBiT and DarkBiT ^10^.

The switch performance is mainly determined by 1) the affinity of the binders, 2) the linker properties which determine effective concentrations within the architecture, and 3) the relative affinities of DarkBiT and SmBiT in the scaffold^10^. As the linker in the dark-LUMABS platform was specifically chosen to span the distance between two epitope binding sites of an antibody, applying the same design strategy to a sensor for multivalent viral targets may prove problematic due to differences in distances between binding sites. Moreover, the integration of multiple binder domains for viral protein targets (instead of simple peptide epitopes) would result in complex, multidomain fusion proteins whose expression could prove difficult due to misfolding during translation. Lastly, designs consisting of linear fusion proteins do not allow the inclusion of binders that require a free N-terminus for target recognition ^17^.

Therefore, the binders and the actuator should be produced separately to avoid these problems. In order to combine binders and actuator into a functional protein switch, we employed the SpyTag/SpyCatcher protein conjugation method developed by the Howarth group ^15^. This method allows to covalently connect two proteins post-translationally via isopeptide bond formation (Fig. 1A) and has been used extensively in applications such as surface modification ^18,19^, fluorescent cell imaging ^20^, and vaccine development ^21^. A highly optimized, third generation of the Spy system has recently been reported ^22^ as well as an orthogonal system named DogTag/DogCatcher, allowing the construction of more complex protein-based architectures ^23,24^. We thus constructed a design named dark-LUX that marries the LUMABS platform with the Spy-technology. The actuator ‘scaffold’ contains both a DarkBiT and a SmBiT that compete for LgBiT complementation and will emit bioluminescence when DarkBiT is spatially separated from LgBiT (Fig. 1B). The DarkBiT is encoded C-terminally of two Catcher domains that are themselves separated by a linker. It is important to note that the scaffold by itself is not responsive to a target as it does not contain any sort of target binding domains. To achieve sensing, the Catcher domains must be functionalized with binders fused to matching tags.

### The dark-LUX design critically depends on intramolecular affinities

We started by expressing three scaffold variants comprising different DarkBiT/SmBiT affinities, named dark-LUX A1 (DarkBiT: 2.5 µM, SmBiT: 190 µM), A2 (DarkBiT: 0.18 µM, SmBiT: 190 µM), and A3 (DarkBiT: 0.18 µM, SmBiT: 2.5 µM). The scaffolds were expressed in *E. coli* and purified via Ni-NTA and StrepTactin affinity purification. As binders, we chose the recently described nanobodies SD36 and SD38 to target IAV HA, as SD36 and SD38 have broad specificity within HA subgroups due to their binding to conserved epitopes at the HA stem region ^25^. SD36 was shown to neutralize IAV group 2 viruses that contain HAs H3, H4, H7, and H10, while SD38 was shown to neutralize some IAV group 2 viruses, but mainly group 1 viruses that contain H1, H2, and H5 ^25^. Both nanobodies were fused to either SpyTag003 and DogTag (4 constructs in total), produced in *E. coli* and purified via StrepTag II and size exclusion chromatography to obtain correctly folded monomers. The formation of branched protein assemblies (Fig. 1A) upon mixing binders with dark-LUX scaffolds was validated via denaturing SDS-PAGE (Fig. 2A). The reaction between SpyTag003/SpyCatcher003 and DogTag/DogCatcher occurred under mild conditions and room temperature within minutes at high nanomolar/low micromolar concentrations. A 1.5-fold molar excess of the binders over the scaffolds was chosen during the assembly reaction to ensure full conversion of the scaffold proteins to functional sensors. Due to the complete conversion, no further purification steps were necessary.

**Figure 2.**
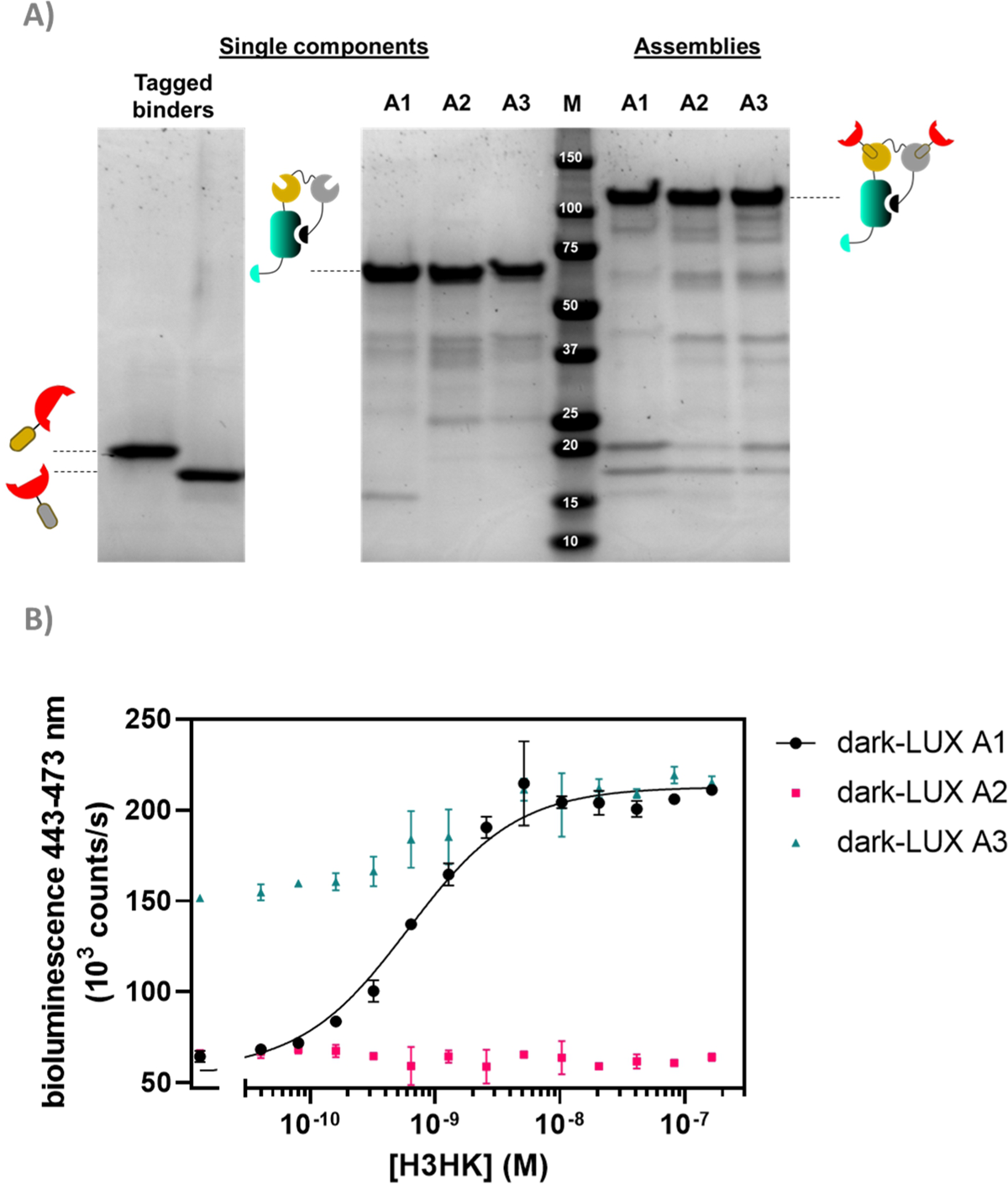
Proof of concept of dark-LUX sensors. A) SDS-PAGE lanes showing the assembly of dark-LUX sensors from single components. Single components: Tagged binders containing tag domains (first lane SD38-Dog, second lane SD38-SpyTag003); A1-A3: respective dark-LUX scaffolds containing Catcher domains; Assemblies: assembled dark-LUX A1, A2 and A3 sensors. B) Titrations for three assembled dark-LUX switches with H3HK. Experimental conditions: Final sensor concentration 200 pM, 1xPBS + 1 mg/ml BSA, final NanoGlo dilution of 1:2000, 2 h incubation at 22 °C. Error bars correspond to the standard deviation of n = 3 technical replicates.

As SD36 was reported to bind H3 with high affinity, ^25^ titration experiments with dark-LUX-SD36 assemblies were performed using trimeric H3 HA from Hongkong (A/Hong Kong/1/1968/H3N2, H3HK) purified from cell culture supernatants ^26^. Dark-LUX A1 showed good luciferase inhibition in the absence of target and was activated upon addition of H3HK, resulting in a 228 ± 12% increase in luminescence intensity with a *K_D,app_* of 613 ± 72 pM and a limit of detection (LoD) of 160 pM (Fig. 2B). Dark-LUX A2 was completely unresponsive and showed a constant level of background bioluminescence, presumably due to the high DarkBiT affinity. Dark-LUX A3 was responsive but showed a high background signal and thus only a modest 42 ± 2% change in bioluminescence, likely caused by inefficient inhibition in the OFF-state due to high SmBiT affinity. Taken together, the data suggest that the proposed sensing mechanism based on bivalent target interaction and subsequent conformational rearrangement works as intended, provided the intramolecular affinities are chosen correctly (Fig. 1B). No change of bioluminescent signal above the background was observed if none or only one SD36 binder was attached to the dark-LUX scaffold (Fig. S1).

### Dark-LUX sensors can detect hemagglutinin variants

After identifying A1 as the optimal scaffold, we next assessed the ability of the platform to sense HAs from different virus groups. Based on *in vitro* neutralization experiments by Laursen *et al.* ^25^, we expected to see a difference in specificity depending on the anti-HA nanobodies used, as SD36 showed better neutralization of IAV group 2 (H3, H4, H7, H10), while SD38 worked better for IAV group 1 (H1, H2, H5, H6) and with less efficiency for some group 2 viruses. Dark-LUX A1 switches with either SD36 or SD38 nanobodies were synthesized and tested on various purified HAs (Fig. 3A). As expected, switches containing SD36 binders showed higher signals on HAs from IAV group 2 compared to SD38 switches, while the latter showed higher signals for most group 1 HAs. Only low levels of unspecific switch activation were observed when a sample with unrelated multivalent protein was used (SARS-CoV2 Spike N-terminal domain (NTD)) ^27^. Titrations were performed with H1 (A/Solomon Islands/3/2006 H1N1, H1Sol) from IAV group 1 for the SD38-based sensor and with H3HK from IAV group 2 for the SD36-based sensor (Fig. 3B,C). The SD38-containing switch bound H1Sol with *K_D,app_* = 1.23 ± 0.15 nM, showing a dynamic range 474 ± 34% and a LoD of 172 pM, while the SD36-containing switch bound H3HK with *K_D,app_* = 0.62 ± 0.06 nM and showed a dynamic range of 543 ± 55% and LoD of 80 pM. Switches that combined one SD36 and one SD38 domain were also responsive and showed binding properties in between those of the SD36- and SD38-only switches (Fig. S2). These results suggest that effective broad HA sensors can be generated by using universal HA binders, but that the specificity of the sensor platform can be tuned towards desired HAs by choosing specific binders.

**Figure 3.**
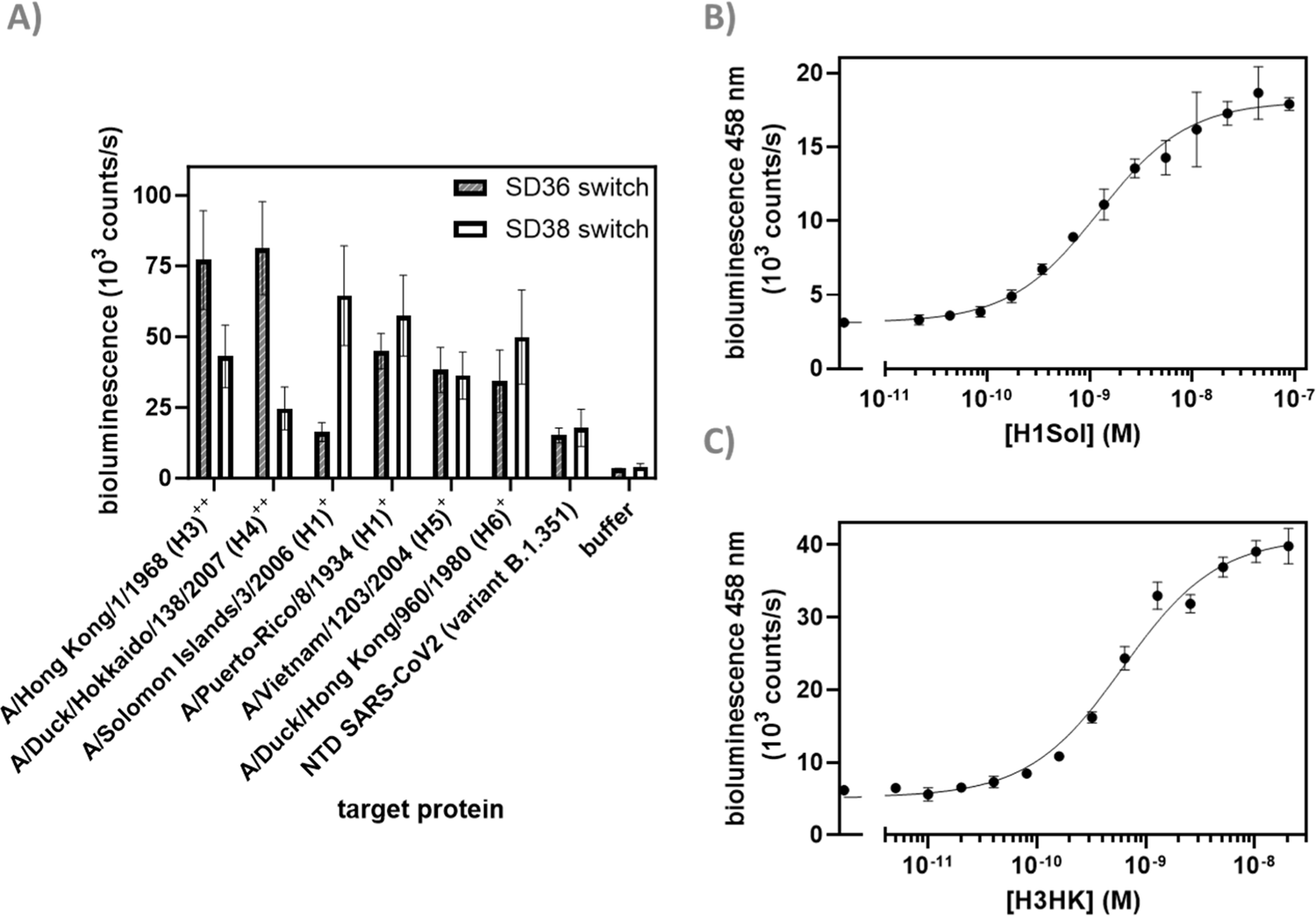
Detection of HA variants. A) Screen of various HA variants from different IAVs. HAs from IAV group 1 and 2 are labeled with ^+^ and ^++^, respectively. Experimental conditions: 250 pM sensor, 8 µg/ml of the respective target proteins or negative control (N-terminal Domain (NTD) SARS-CoV-2), incubation 2 h 22 °C; B) Titration of H1Sol with SD38 dark-LUX. 200 pM sensor, incubation 16 h at 4 °C; C) Titration of H3HK with SD36 dark-LUX. 150 pM sensor, incubation 16 h 4 °C; Further experimental conditions were the same in all experiments: 1xPBS + 1mg/ml BSA, final NanoGlo dilution of 1:2000. All error bars correspond to the standard deviation of n = 3 technical replicates.

### Dark-LUX sensors can detect N2 neuraminidase

To further broaden the scope of the sensor platform, we next targeted neuraminidase (NA) as an additional relevant target on the surface of IAVs using a different type of binder, single chain variable fragments (scFvs). scFvs combine the advantages of antibodies (high specificity and affinity) while being considerably smaller and easier to express. Thus, we constructed a scFv binder from an N2-specific antibody AS4C that was identified by the group of Alain Townsend (unpublished) and expressed it with a DogTag in *Pichia pastoris* to achieve properly folded protein. Microscale thermophoresis (MST) experiments with Alexa647-labeled AS4C-DogTag demonstrated good binding to N2 (A/NL/354/16, N2-16, *K_D,app_* ≈1 nM) (Fig. S3). As expression of the scFv with SpyTag003 was not successful, we exchanged the SpyCatcher003 in the dark-LUX A1 with a second DogCatcher domain and assembled a switch with two AS4C scFvs on this scaffold for titrations against N2-16. An increase in bioluminescence signal was observed upon the addition of N2-16 (dynamic range 137 ± 13 %), yielding a *K_D,app_* of 3.87 ± 0.75 nM (Fig. 4). The sensor did not respond if an N9-type neuraminidase was used as a target, in line with the N2-specificity of AS4C (Fig. S4). While both dynamic range and sensitivity were lower compared to the HA sensors, the N2-16 dark-LUX sensor clearly switched from a closed to an open conformation. Importantly, dark-LUX sensors that were functionalized with either HA or NA binders selectively detected their respective target and did not show cross-reactivity (Fig. S5).

**Figure 4.**
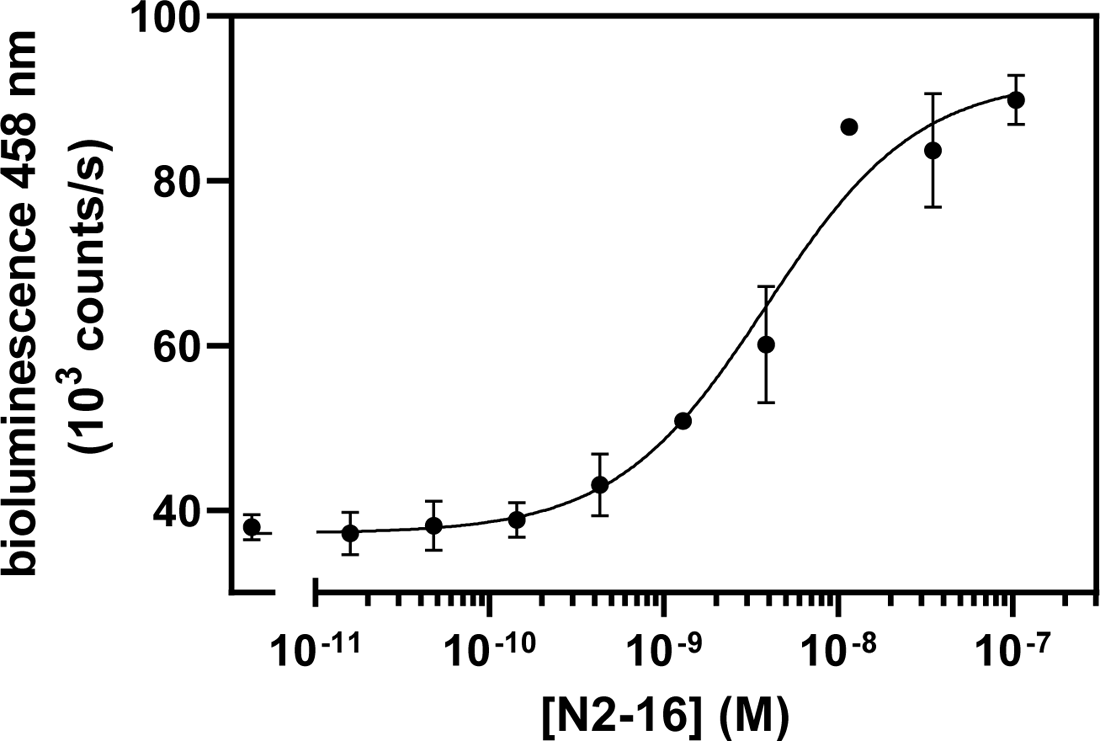
Titration on A/NL/354/16 (N2-16) with AS4C scFv binders. 200 pM sensor, incubation 16 h at 4 °C. Further experimental conditions: 1xPBS + 1mg/ml BSA, final NanoGlo dilution of 1:2000. All error bars correspond to the standard deviation of n = 3 technical replicates.

### Linker choice in homobivalent dark-LUX sensors is less critical than expected

In previously published works on fusion protein switches, changes in linker lengths and identity were key to obtain variants with high signal-to-noise ratio ^16,28^. This is because linkers play critical roles in providing flexibility, mediating conformational changes in the tethered domains that are required for switch activation, and spanning the distance between the targeted epitopes/receptors ^29,30^. To allow efficient switching of the dark-LUX sensors, the linkers that connect the domains of the initial design were carefully chosen. Both Catchers are connected via a long, flexible poly-GS linker (61 aa) with the rationale to provide sufficient flexibility for the Catchers to be separated in the target bound (ON-) state, while still allowing DarkBiT to inhibit LgBiT in the target unbound (OFF-) state. On the other hand, LgBiT and DarkBiT are fused with relatively short linkers to DogCatcher (6 aa) and SpyCatcher003 (9 aa), respectively, to ensure their separation in the ON-state conformation.

To investigate the influence of linker properties on the performance of dark-LUX sensors, a plasmid library was constructed to create, express, and sample variants of the dark-LUX A1 design using the iFLinkC technology. This cloning technology relies on iterative recombination of separately cloned *functional domain* and *linker* plasmids using type II S restriction enzymes and allows to simultaneously vary all linker positions (L1, L2, L3) ^16^. We chose to sample short linkers (2-9 aa) at positions L1 and L3 and longer linkers (15-47 aa) at position L2 to maintain the initial short-long-short linker combination that allows sensor switching (Fig. 1, Table S1). After library assembly and transformation of *E. coli* BL21, scaffold variants from ∼100 single colonies were expressed in deep well plates. Following lysis, scaffolds were batch-purified using StrepTactin plates, sensors assembled by the addition of tagged SD38 binders, incubated with H1Sol or buffer, and screened to determine variants that showed increased luminescent signal in the presence of target. We purified selected scaffold variants that all had different linkers for L1, L2, and L3 as confirmed by sequencing (Table S2). Despite this variety in linker sequences, the tested variants showed overall similar sensor properties compared to the original A1 sensor when used in titrations (Fig. 5A), with *K_D,app_* and dynamic range values in the same order of magnitude as the original design. These results indicate that the exact length and constituency of linkers in the dark-LUX scaffold is less critical. The possibility to screen linker lengths and constituency remains an important feature of the dark-LUX platform design, however, in particular when considering other multivalent targets where larger distances must be spanned^31^.

**Figure 5.**
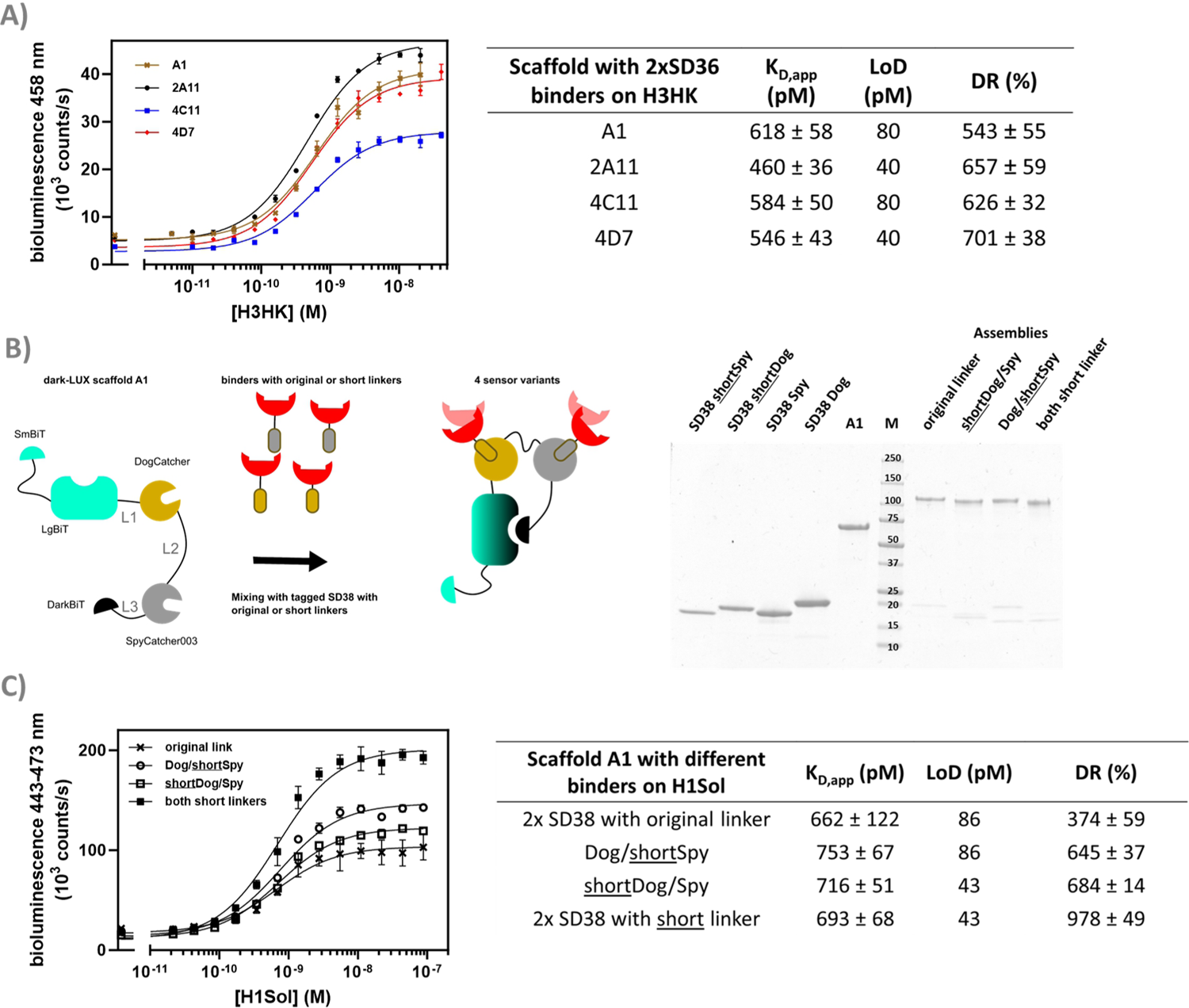
Linker screening and optimization. A) Selected variants of the iFLinkC based linker library of dark-LUX A1 assembled with S36 binders were titrated against H3HK. B) Assembly schematic of sensor variants used to investigate binders with shorter linkers. The SDS-PAGE shows how dark-LUX scaffold A1 (lane A1) was assembled with tagged SD38 variants into functional dark-LUX sensors. SD38 variants include short linkers (SD38 shortSpy and SD38 shortDog) or original ones (SD38 Spy, SD38 Dog) C) H1 Solomon Island titrations with assembled sensors from B) containing SD38 binders with different linkers show differences in dynamic range. Experimental conditions: Final sensor concentration 200 pM, 1xPBS + 1mg/ml BSA, final NanoGlo dilution of 1:2000, 16 h incubation at 4 °C. Error bars correspond to the standard deviation of n = 3 technical replicates.

Aside from linkers in the scaffold, 19 aa linkers were originally introduced between the binders and the DogTag/SpyTag003 to ensure efficient conjugation and allow interactions between sensor and target. However, if these linkers are too long, this may prevent full dissociation of DarkBiT and LgBiT in the ON-state. To test this hypothesis, the original linkers between SD38 and the DogTag/SpyTag003 were shortened to 10 aa. Dark-LUX sensors were assembled that contained SD38 binders with the original 19 aa linkers or shorter ones (Fig. 5B) and subsequently used in titrations against H1Sol. While the effect on the *K_D,app_* was not significant, the dynamic range increased due to a higher activity in the target-bound state, which also translated into higher sensitivity for scaffolds functionalized with SD38-DogTag that had a shortened linker (LoD: 43 pM) instead of the longer one (LoD: 86 pM) (Fig. 5C). These results suggest that shorter linkers between DogTag/SpyTag003 and the binder lead to better separation of DarkBiT from LgBiT in the target-bound conformation (ON-state) of the switch.

### A heterobivalent dark-LUX sensor shows pronounced linker dependency

Discriminating between influenza strains is important for epidemiology and surveillance ^32^. We reasoned that the modular dark-LUX platform should allow the detection of a certain HA variant by combining a binder specific to the variable head region of a given HA with a broadly binding stem binder. This is possible with the dark-LUX scaffold as it contains two different Catcher domains, allowing to specifically introduce two different binders. Recently, advances in computational protein design have resulted in *de novo* binders with high affinity and specificity to their targets ^33,34^. For example, the small binder HSB2.A (9.5 kDa, 86 aa) was computationally designed to bind H3HK (reported *K_D_* ≈ 0.5 nM) ^34^. Therefore, we functionalized the dark-LUX A1 scaffold with HSB2.A-DogTag as ‘head binder’ and SD36-SpyTag003 as ‘stem binder’ and tested the assembled switch in titrations on purified H3HK. The sensor worked and was specific, as a control where SD36 was exchanged to SD38 and a control with H1 from A/Puerto-Rico/8-1934 as target both showed only minimal sensor response (Fig. S6). The sensor showed a relatively modest dynamic range of 63 ± 18 % (LoD: 5.1 nM) compared to sensors with two stem binders (Fig. 5A, Fig. 6A), which may be caused by a different distance between binding sites. Estimations based on crystal structures showed that the distance between two SD36 stem binding sites is approximately 50 Å, while the distance between the head region of H3HK and its corresponding stem binding site is considerably larger with 80 Å (Fig. 6B). To check whether a more rigid linker at position L2 would improve the performance of the head/stem binding switch ^13,14,35^, we exchanged the flexible GS-rich linker in the A1 scaffold for (TP)_n_ linkers. This naturally occurring linker motif has properties of rigid linkers ^36^, arguably due to high proline content that causes higher persistence lengths ^37^. When used in titrations on purified H3HK, dark-LUX scaffold variants with (TP)_15_ and (TP)_22_T linkers indeed performed better than the original A1 sensor and yielded a dynamic range of 110 ± 9 % (LoD: 1.3 nM) for the (TP)_22_T variant and 124 ± 7 % (LoD: 321 pM) for the (TP)_15_ variant (Fig. 6A). All sensors showed comparable *K_D,app_* values between 1.2 and 1.4 nM for H3HK binding.

**Figure 6.**
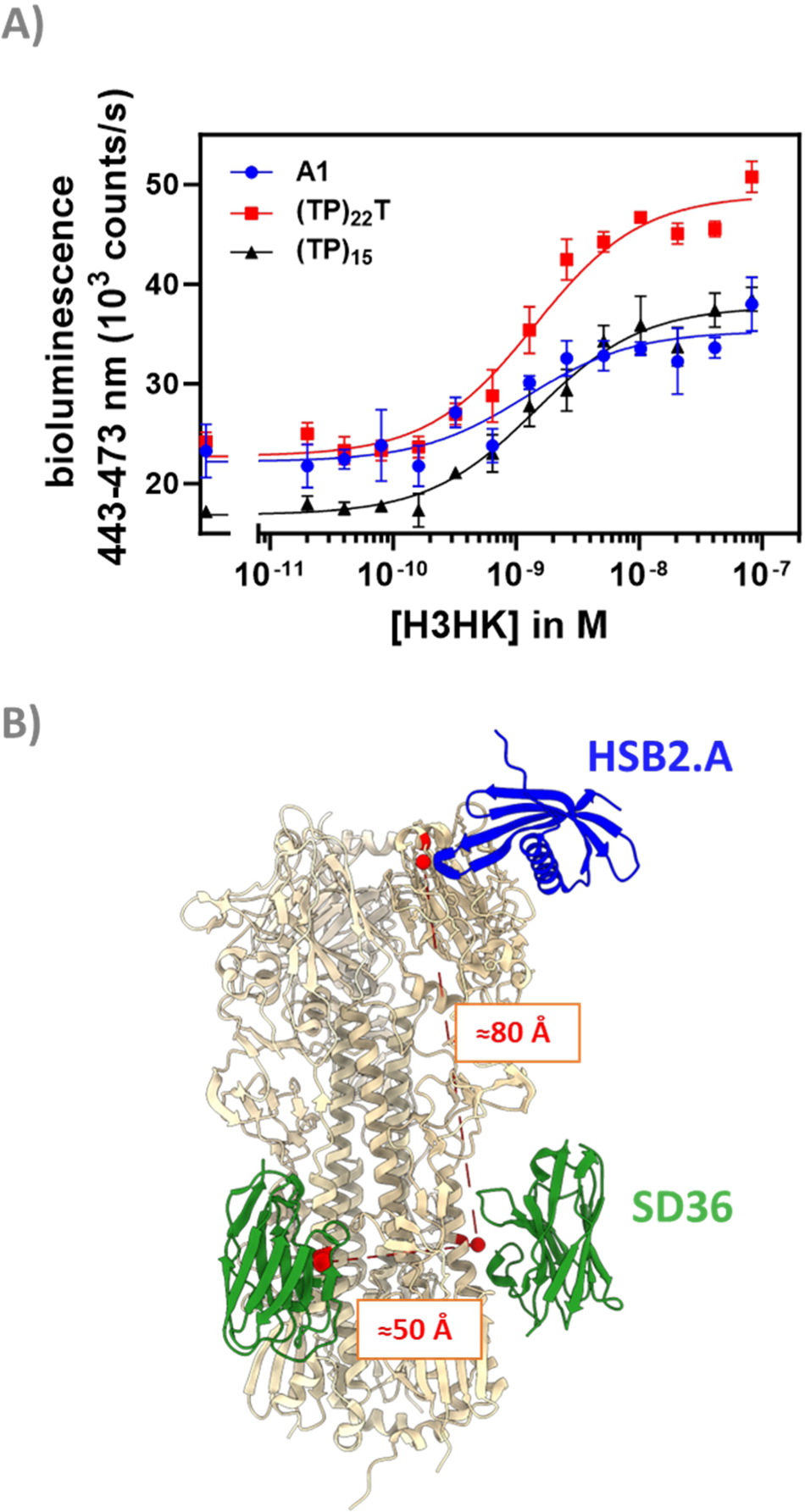
Performance and conceptualization of head & stem binding dark-LUX switches. A) H3HK titrations for the tested dark-LUX switches A1 (containing 58 aa GS-rich linker), (TP)_22_T and (TP)_15_. Experimental conditions: Final sensor concentration 300 pM, 1xPBS + 1mg/ml BSA, final NanoGlo dilution of 1:2000, 16 h incubation at 4 °C. Error bars correspond to the standard deviation of n = 3 technical replicates. B) Distances between head and stem binding sites of hemagglutinins (shown in red) were estimated based on PDB crystal structures 5KUY and 6FYU. Structures visualized with UCSF Chimera ^44^.

## Conclusions

The herein presented sensor strategy harnesses two features to construct modular, branched proteins that can detect targets with a high sensitivity. Firstly, the complementation-inhibited NanoBiT scaffold accounts for a low background bioluminescent signal in the absence of target molecules while producing bright blue light when conformationally switched, provided the intramolecular affinities of the complementation partners are well adjusted to allow switching. Secondly, the SpyCatcher/SpyTag technology allows fast, covalent conjugation of different binders to the scaffold, representing a highly reliable bioconjugation strategy. To understand how sensitivity and sensor activation are affected, we screened various parameters of the platform including 1) the binding domains, 2) the intramolecular affinities of DarkBiT and SmBiT, 3) the linkers between the scaffold components, and 4) the linkers between the binders and the SpyTag003 / DogTag. All classes of binders tested in this work, i.e. nanobodies (SD36, SD38), computationally designed binders (HSB2.A) and scFvs (AS4C), could be conjugated to the scaffold platform, and titrations with these assemblies allowed bioluminescent detection of pM-nM concentrations of viral surface proteins.

The platform described here is well-suited to enable straightforward introduction of novel binders and rapid optimization of bioluminescent sensor properties. The rapid development of binders for novel biomolecular targets becomes more and more feasible, both as result of deep learning algorithms for de novo protein design and the introduction of more efficient methods for directed evolution^41–43^. E.g. during surveillance of poultry farms specific detection of IAV H5 and H7 subtypes can be used to determine whether outbreaks are caused by virus variants of concern. This could be achieved by functionalization of dark-LUX sensors with newly developed binders that target the multibasic cleavage site, a unique H5 and H7 motif of highly pathogenic avian IAV ^40^.

To arrive at sensitive viral antigen tests that at the same time fulfill the main POC requirements (fast, simple-to-use, homogeneous detection), the multivalent presentation of viral surface proteins can be exploited by constructing biosensors that bind to multiple, potentially different target proteins on the virus surface, which would allow a more specific detection of virus variants depending on the targeted HA and NA subtypes. Systems with higher valency could also be constructed by including multiple SpyCatcher and DogCatcher domains in the scaffold and mixing them with binders ^39^. These sensors might even allow the direct detection of intact, infectious virus using simple readout in a homogeneous test at the point-of-care.

## Supporting information

Supplement Grawe et al

## Author contributions

Alexander Gräwe: Conceptualization, Methodology, Investigation, Writing – original draft, Writing – review & editing; Cindy M. Spruit: Resources (supporting), Writing – review & editing. Robert P. de Vries: Conceptualization (supporting), Writing – review & editing; Maarten Merkx: Conceptualization, Methodology, Writing – original draft, Writing – review & editing.

## Acknowledgements

We would like to thank Pramila Rijal and Alain Townsend for fruitful discussions and for kindly providing the sequence of the N2-specific AS4C antibody. We would like to thank the members of the Merkx group for fruitful discussions, in particular Anna Świetlikowska, Tallie Godschalk, Jora Couwenberg, and Imke Goertz. This project has received funding from the European Union’s Horizon 2020 research and innovation programme under the Marie Skłodowska-Curie grant agreement No 899987.

