## Supplement Grawe et al for "Bioluminescent detection of viral surface proteins using branched multivalent protein switches"

**Table of contents**

Supplementary Materials and Methods S2
- Materials S2
- Cloning S2
- Protein purification of Dark-LUX variants and binders S2
- Protein purification of viral surface proteins S3
- Sensor assembly S3
- Library sampling S3
- Titrations S4
- MST experiments S4

Supplementary Tables and Figures S6
- Table S1: Overview of linker library S6
- Table S2: Sequence-verified scaffold variants S6
- Figure S1: dark-LUX switch controls S7
- Figure S2: Titrations with SD36 and SD38 combinations S7
- Figure S3: Microscale Thermophoresis of AS4C binding to N2 S8
- Figure S4: Selectivity test of N2 sensor S8
- Figure S5: Selectivity test of HA and NA sensor S9
- Figure S6: Selectivity test of sensor with head/stem binder S10

Protein sequences S11

References S14

**Materials**

Unless stated otherwise, chemicals were purchased from Sigma or Carl Roth.

**Cloning**

DNA nucleotides and gblocks were purchased from IDT, and cloning and plasmid amplification was performed in *Escherichia coli* TOP10. To obtain initial dark-LUX scaffold A1, the gblock was inserted into pET28a(+)_dark-LUMABS CTX using Kpn1-HF (NEB) & Spe1-HF (NEB) restriction and ligation. To obtain scaffold A2 and A3, SmBiT and DarkBiTs were exchanged for variants with different affinities using restriction enzymes and T4 ligase (NEB) to combine parts from previously described pET28a(+)_dark-LUMABS plasmids with dark-LUX inserts.

gBlocks containing binders (SD36, SD38, HSB2.A) were cloned into pET28 using similar methods. For AS4C-HL-DogTag (HL: heavy-light conformation of the scFv), the gblock was inserted into pPICZalphaB using pPICZalphaB-SapL3 (addgene plasmid #78171) as template and Pst1-HF & Not1-HF restriction and T4 ligation. All constructs were verified via Sanger Sequencing (Azenta Life Sciences).

pPICZalphaB_AS4C-HL-DogTag was integrated into the genome of *Pichia pastoris* X33 following the EasySelect protocol (Invitrogen) [1]. Briefly, plasmid DNA was linearized with Sac1-HF and concentrated on an Amicon Ultra-4 (Merck Millipore) with 10 kDa MWCO cutoff to remove the buffer and dissolve the DNA in sterile water. The retentate was transferred to a 0.5 ml Amicon with 10 kDa MWCO cutoff for further concentration of DNA. 80 µl of *P. pastoris* cells from a fresh culture were mixed in an electroporation cuvette with 10 µl of purified & concentrated DNA and electroporated with 1.5 kV and 5.6 ms pulse time. Cells were incubated at 2.5 h at 30 °C and afterwards plated on YPDS plates supplemented with Zeocin. Plates were incubated until colonies formed. Colonies were checked for successful integration with colony PCR according to the manual [1].

The linker library was generated using the previously described iFLinkC method [2]. The final library was cloned into pFLinkC-XE to allow screening via IPTG induced expression in *E. coli* BL21 (DE3) cells. Linkers used in the library are listed in Table S1.

**Protein purification of Dark-LUX variants and binders**

For purification of dark-LUX variants and Spy/Dog-tagged binders (except AS4C- HL-DogTag), *E. coli* BL21 (DE3) cells were transformed with corresponding pET28a(+) plasmids. Cells were grown in LB medium supplemented with 50 µg/ml kanamycin at 37 °C at 180 rpm in a shaking incubator. Large cultures (0.5-2 l) in 2 l or 5 l baffled flasks were inoculated with corresponding overnight cultures and induced with IPTG at $OD_{600}$ 0.6-0.8. Proteins were expressed overnight at 18 °C. Harvested cells were lysed with BugBuster reagent (Novagen) supplemented with Benzonase (Merck), and proteins were purified according to their tags (see sequences of used proteins) with Ni-NTA chromatography and/or Strep-Tactin XT (iba) using gravity flow columns. For SD36 and SD38 variants, an additional size exclusion step was performed to achieve nanobody monomers, using 1xPBS (10 mM phosphate buffer, 2.7 mM KCl, and 137 mM NaCl, pH 7.4) as running buffer on a HighLoad 16/600 Superdex 75pg column.

Protein purity was confirmed by reducing SDS-PAGE and concentrations were calculated using $A_{280}$ Nanodrop measurements using the corresponding theoretical extinction coefficients (based on protein sequence). Proteins were aliquoted, flash frozen in liquid $N_2$ and stored at -70 °C.

For purification of AS4C-HL-DogTag, *P. pastoris* cells with genomically integrated AS4C-HL-DogTag were grown in BMGY medium and transferred to BMMY medium for induction of the AOX promoter with MeOH. The protein was present in the medium 24 h after induction as observed via SDS-PAGE. Medium was harvested after an additional 2 d, spun at 40000xg, 4 ˚C for 30 min and supernatant was applied to equilibrated Ni-NTA columns for purification of the secreted AS4C-HL-DogTag via the C-terminal 6xHisTag. Elutions were buffer exchanged to 1xPBS via PD10 columns (Cytiva), flash frozen, and stored at -70 °C.

**Protein purification of viral surface proteins**

Cloning of trimeric HAs and tetrameric NAs was performed as described previously [3]. Shortly, codon-optimized NA genes (A/NL/354/16 [EPI_ISL_355168] (N2) and A/Sh/02/13 [YP_009118481] (N9) were cloned into a mammalian expression vector, preceded by sequences successively coding a StrepTag II and a TB tetramerization motif. The used expression vector is addgene plasmid #158742 [3], with the removal of the sfGFP. Codon optimized genes coding for hemagglutinins (HA) from A/Solomon Islands/3/2006 (H1N1), A/Hong Kong/1/1968 (H3N2), A/Duck/Hokkaido/138/2007 (H4N6), A/Vietnam/1203/2004 (H5N1), A/Duck/Hong Kong/960/1980 (H6N2) [4], A/Puerto-Rico/8/1934 (H1N1) were cloned in an expression vector (addgene plasmid #157933 [5]) that codes for a StrepTag II and a GCN4 trimerization domain in frame with the HA gene, with the removal of the sfGFP. NAs and HAs were purified from cell culture supernatants after expression in HEK293T or HEK293S GnTI(-) cells, respectively, as described previously [6]. In short, transfection was performed using the pCD5 expression vectors and polyethyleneimine I, in a ratio of 1:8 for HA and 1:12 for NA. The transfection mixtures were replaced at 6 h post-transfection by 293 SFM II expression medium (Gibco), supplemented with sodium bicarbonate (3.7 g/L), Primatone RL-UF (3.0 g/L, Kerry, NY, USA), glucose (2.0 g/L), glutaMAX (1%, Gibco), valproic acid (0.4 g/L) and DMSO (1.5%). At 5 to 6 days after transfection, tissue culture supernatants were collected and Strep-Tactin sepharose beads (iba, Germany) were used to purify the HA proteins according to the manufacturer's instructions.

**Sensor assembly**

For each assembly reaction, a purified Dark-LUX scaffold containing DogCatcher/SpyCatcher003 was mixed with two binders containing DogTag/SpyTag003 at a molar ratio of 1:1.5:1.5 to guarantee complete conversion of all added scaffold into functional sensor. The mix was shortly spun down and incubated overnight at 4 °C. The scaffold concentration was generally 2 µM but at least 500 nM to allow complete isopeptide bond formation for the DogTag/DogCatcher system after overnight incubation [7]. Reactions were performed in a final volume of 40 µl and with 1xPBS as buffer.

**Library sampling**

Single colonies of *E. coli* BL21 (DE3) cells transformed with the pFLinkC-XE dark-LUX scaffold library were used to inoculate 750 µl LB plus 100 µg/ml ampicillin (+Amp) each in a 96-well deepwell plate. Cells were grown overnight at 37 °C at 250 rpm and used to inoculate 750 µl LB+Amp to an approximate OD of 0.1. After 3 h of additional growth, IPTG was added to a final concentration of 1 mM, and expression was conducted for 3.5 h at 37 °C. Cells in the deepwell plate were harvested by centrifugation at 2500xg, 4 °C, 15 min, and cell pellets stored at -20 °C overnight. Cells were resuspended in 200 µl of lysis buffer (150 mM NaCl, 100 mM Tris-Cl pH=8, 1 mg/ml lysozyme (Roth), 2 u/ml DNase I (NEB)) and lysed for 3 h at 30 °C and 300 rpm. After centrifugation at 2500xg, 4 °C, 20 min, supernatants were transferred to a 96 well plate, and expressed dark-LUX scaffolds were purified in parallel using 96-well StrepTactin plate (iba) with two washing steps of 300 µl buffer W (150 mM NaCl, 100 mM Tris-Cl pH=8). Proteins were eluted with buffer E (150 mM NaCl, 100 mM Tris-Cl pH=8, 50 mM biotin). Dark-LUX sensors were assembled by the addition of SD38-DogTag and SD38-SpyTag003 in a final volume of 100 µl with a final concentration of tagged binders of 60-90 nM at room temperature for 2 h. The assembled sensors were mixed with 2 µg/ml H1 Solomon Island (final concentration) or buffer, incubated for 20 min at RT, before furimazine was added to a final dilution of 1:1000 and intensity of bioluminescent signals were recorded using a TECAN Spark 10 M at a final assay volume of 50 µl.

### Titrations

Titrations were performed as described previously in [8]. Briefly, all titrations were performed in 1xPBS supplemented with 1 mg/ml BSA to avoid unspecific adsorption of proteins to surfaces. For incubation at 4 °C overnight, low protein binding tubes were additionally used to prevent adsorption. NanoGlo substrate (furimazine, Promega) was added from a freshly prepared 10x stock directly before the measurement to obtain the final volume and concentrations. Unless stated otherwise, the final NanoGlo dilution during the measurements was 1:2000. Binding assays were performed in white 364-well plates (flat bottom, Nunc or Greiner) and a total volume of 20 µl with n=3 technical replicates. Unless stated otherwise, shown bioluminescence values were determined from bioluminescence spectra (398-653 nm, integration time 100 ms) measured at 22 °C in a TECAN Spark 10M. The resulting binding curves were fitted to the standard binding model (P * x) / ($K_D$ + x) + c, with x being the target concentration and c the background signal. Dynamic range (DR) was calculated via

$$\mathrm{DR} = \frac{\mathrm{Max} - \mathrm{Min}}{\mathrm{Min}}$$

with *Max* being the signal at saturated sensor (mean of n=3) and *Min* the signal in absence of the target (mean of n=3). DR uncertainties were propagated from the standard deviation of *Max* and *Min* (*s(Max)* and *s(Min)*, respectively) using [9]

$$\frac{1}{Min^2}\sqrt{(Max \cdot s(Min))^2 + (Min \cdot s(Max))^2}$$

The limit of detection (LoD) of each sensor was calculated by applying the 3σ rule (mean of blank plus 3 times its standard deviation) as in previous studies [8].

### MST Experiments

To estimate the $K_D$ of the constructed AS4C-HL-DogTag scFv for binding to N2 Virginia, purified AS4C-HL-DogTag was labeled with Alexa647 and used in MST binding experiments. For labelling, the protein was concentrated using an Amicon filter (10 MWCO) and buffer exchanged to 0.2 M sodium bicarbonate pH 8.3 using PD SpinTrap G-25 columns (Cytiva) according to the manufacturer’s protocol. 100 µl of the protein was mixed with 10 µl of Alexa-647 NHS Ester (Lumiprobe) freshly dissolved in DMSO (10 mg/ml) and incubated for 1 h at

22 °C with constant shaking. The reaction was purified from excess dyes by subsequently applying it to PD SpinTrap G-25 column twice. The concentration of the protein and the efficiency of labeling was calculated according to [10]. Labelled AS4C-HL-DogTag (final concentration 500 pM) was mixed with varying concentrations of N2 Virginia and incubated for 1 h at 22 °C in a volume of 40 µl in 1xPBS & 0.02% Tween20. Capillaries were loaded and MST experiment was performed at 60% LED power and 40% MST power on a Monolith NT.115 (Nanotemper Technologies). Two independent experiments were performed, once with standard capillaries (MO-K022) and once with premium capillaries (MO-K025).

Table S1 Overview of linkers included to generate a dark-LUX scaffold library.

| LgBiT - **Linker 1** - DogCatcher | DogCatcher-**Linker 2**-SpyCatcher003 | SpyCatcher003-**Linker 3**-DarkBiT |
|---|---|---|
| GG | G(TP)$_{20}$G | GG |
| GGG | G(TP)$_{25}$G | GGG |
| GGGG | G(TP)$_{30}$G | GGGG |
| GGSG | G(TP)$_{35}$G | GGSG |
| GPPPPG | G(TP)$_{40}$G | GPPPPG |
| GPPPPPPPG | G(TP)$_{45}$G | GPPPPPPPG |
| GPPPG | G(KPPP)$_{5}$KG | GPPPG |
| GGASPAGG | G(KPPP)$_{8}$KG | GGASPAGG |
| GGSGGSG | G(KPPP)$_{10}$KG | GGSGGSG |
| | GGSGGSGGSGGSGSG | |
| | G(P)$_{19}$G | |
| | GA(EAAAK)$_{2}$AG | |
| | G(AEAAAKAGS)$_{3}$G | |
| | GP$_{18}$(TP)$_{7}$G | |

Table S2 Sequence-verified scaffold variants obtained via linker library screening. The change ON/OFF in library screening refers to the change of bioluminescence from the OFF state (buffer added) to the ON state (target added).

| variant | L1 | L2 | L3 | performance (change ON/OFF in library screening) | performance (dynamic range after purification) |
|---|---|---|---|---|---|
| A1 | SSG GGT | KL(GGS)$_{8}$GG ENLYFQS(G GS)$_{7}$GTG | TSGGSS GGS | 320% | 543 ± 55 % |
| 2A11 | GG | G(TP)$_{20}$G | GPPPG | 549% | 657 ± 59 % |
| 2B2 | GPP PG | G(TP)$_{20}$G | GGSGGS G | 270% | n.a. |
| 2B3 | GGA SPA GG | G(TP)$_{45}$G | GPPPG | 420% | n.a. |
| 2B11 | GGS G | G(TP)$_{35}$G | GGASPA GG | 455% | n.a. |
| 4C11 | GPP PPPP PG | G(TP)$_{20}$G | GGASPA GG | 197% | 626 ± 32 % |
| 4D7 | GPP PG | G(TP)$_{30}$G | GPPPPPP PG | 213% | 701 ± 38 % |
| 1KPPP | GGA SPA GG | G(KPPP)$_{10}$K G | GPPPG | 119% | 387 ± 31 % |

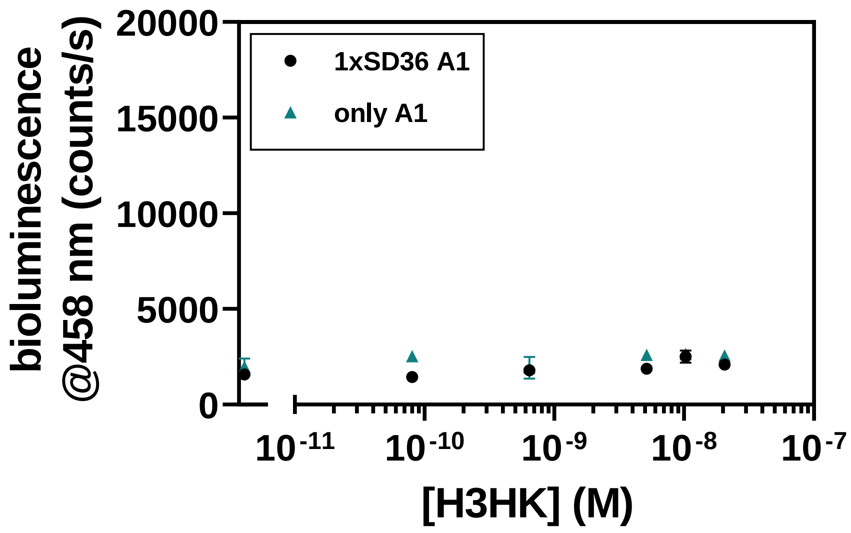


Figure S 1: Controls with empty scaffold, or a scaffold to which only one binder (here: SD36-SpyTag) was attached, do not show an increase in bioluminescence at higher concentrations of target (here: H3 HA from A/Hong Kong/1/1968/H3N2). Experimental conditions: Final sensor concentration 150 pM, 1xPBS + 1mg/ml BSA, final NanoGlo dilution of 1:1000, 16 h incubation at 4 °C. Error bars correspond to the standard deviation of n = 3 technical replicates.

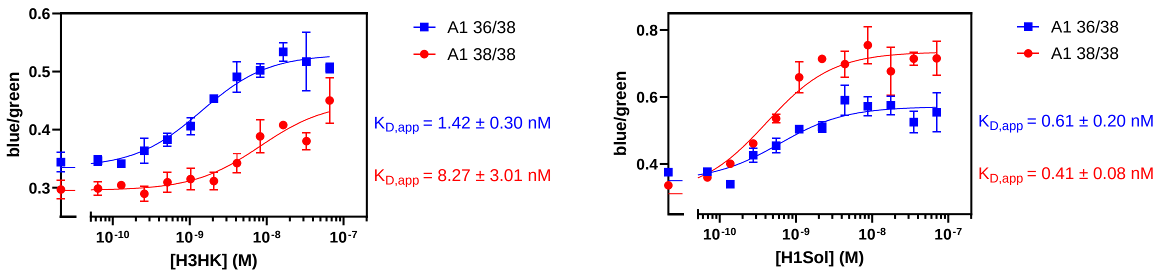


Figure S 2: Titrations with dark-LUX switches containing each one SD36 and SD38 nanobody compared to switches with 2x SD38 nanobodies on H3HK and H1Sol targets. Dynamic ranges and $K_{D,app}$ values change when different binders are used. Experimental conditions: Final sensor concentration 250 pM, 1xPBS + 1mg/ml BSA, final NanoGlo dilution of 1:2000, 2 h incubation at 22 °C. Error bars correspond to the standard deviation of n = 3 technical replicates. For these experiments, the bioluminescence at 458 nm (blue) was normalized to the bioluminescence at 518 nm (green) generated by a calibrator luciferease that was added at a final concentration of 1 pM [8].

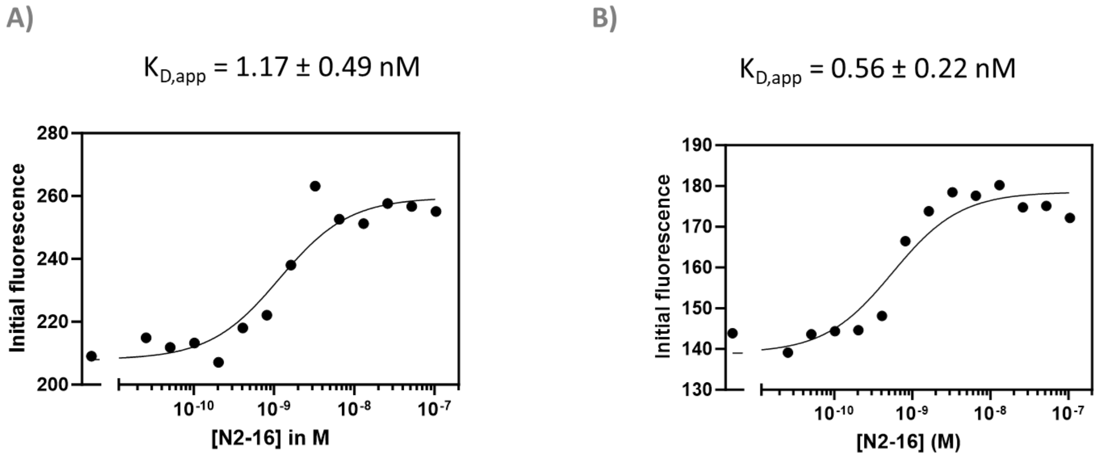


Figure S 3: Microscale thermophoresis (MST) experiments to determine a $K_{D,app}$ for AS4C HL binding to N2-16. The experiment was performed twice, each time with different capillaries. 1xPBS + 0.02% Tween20 was used as buffer. A) standard capillaries, 1 nM labelled AS4C HL B) premium capillaries, 500 pM labelled AS4C HL. The mixtures were incubated 1 h at 22 °C (room temperature) before they were transferred into capillaries and measured.

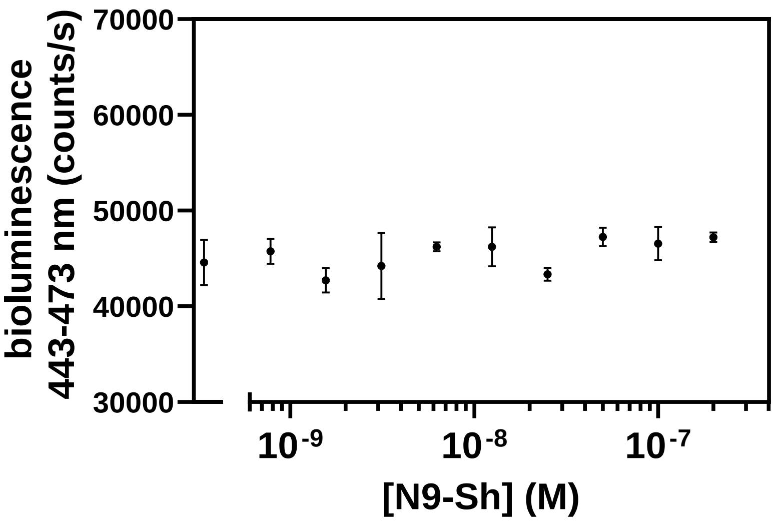


Figure S 4: Selectivity test of N2 sensor (dark-LUX A1 scaffold functionalized with 2x AS4C HL scFv) on a different neuraminidase target, N9 from A/Sh/02/13 (N9-Sh). The sensor does not respond to N9-Sh due to the N2-specificity of AS4C HL. Experimental conditions: Final sensor concentration 300 pM, 1xPBS + 1mg/ml BSA, final NanoGlo dilution of 1:2000, 16 h incubation at 4 °C. All error bars correspond to the standard deviation of n = 3 technical replicates.

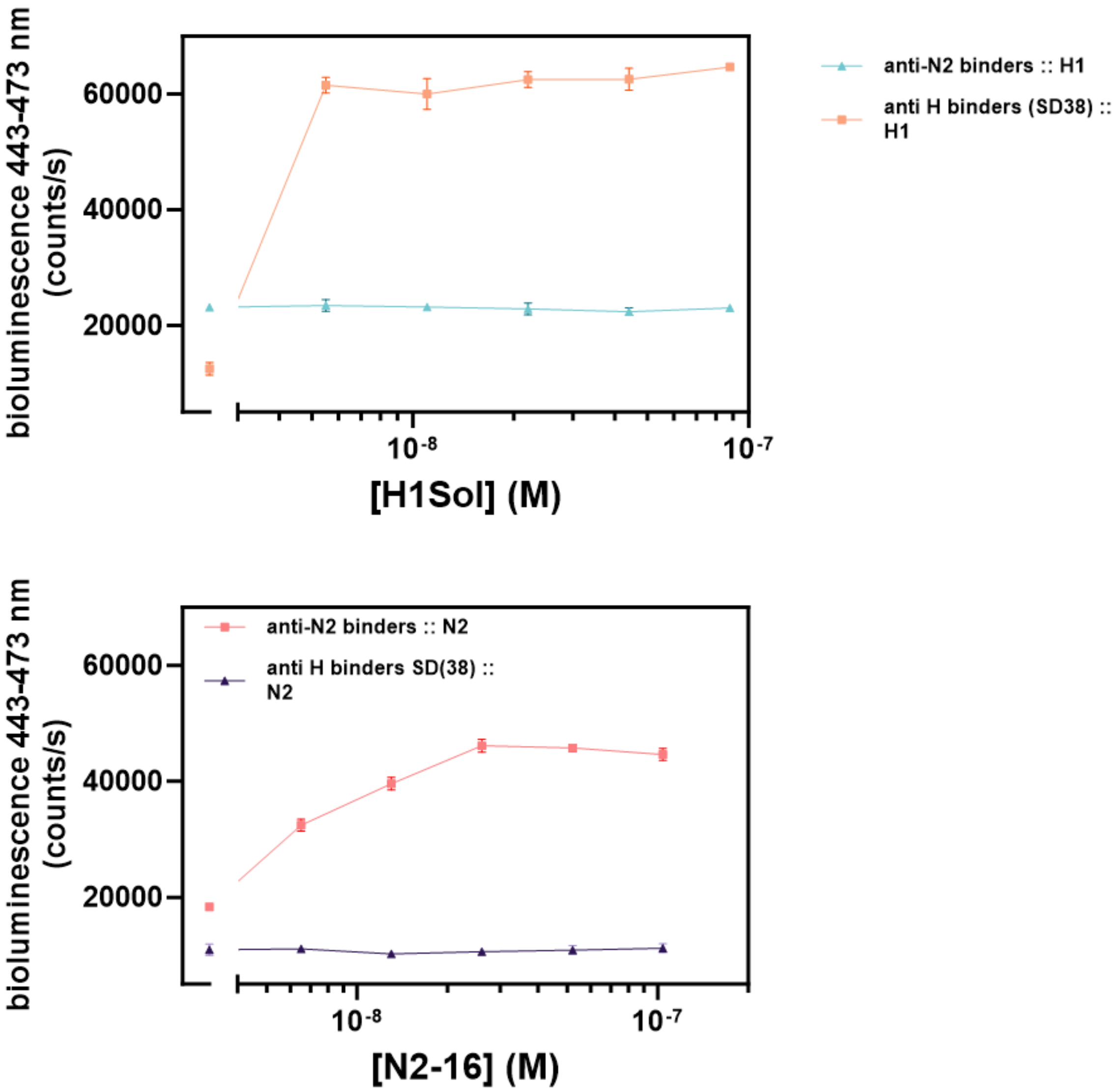


Figure S 5: Selectivity tests using Dark-LUX A1 with 2xSD38 binders and Dark-LUX A1 with AS4C-HL binders. Experimental conditions: Final sensor concentration 300 pM, 1xPBS + 1mg/ml BSA, final NanoGlo dilution of 1:2000, 1 h incubation at 22 °C. Error bars correspond to the standard deviation of n = 3 technical replicates. Top: H1Sol as target, Bottom: N2-16 as target. The respective sensors are activated only in presence of the correct target.

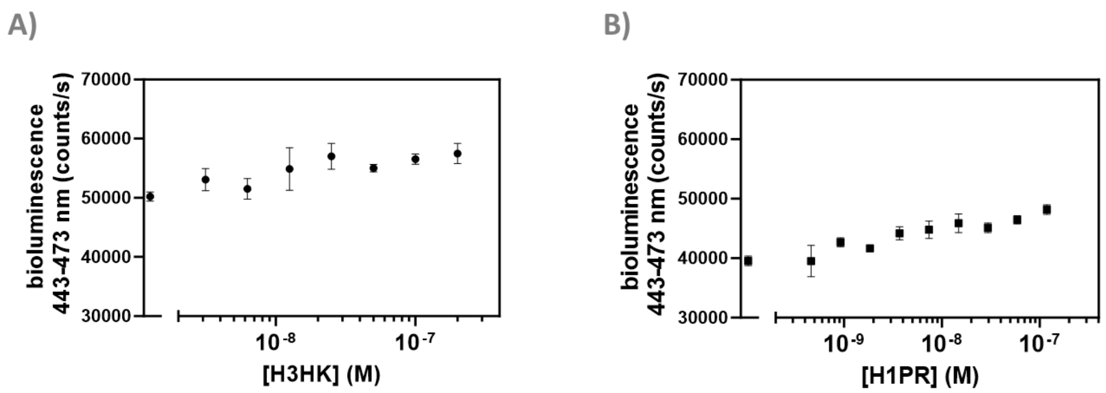


Figure S 6 Selectivity tests with HSB2.A & stem binder switches. A) Titration on H3HK with a dark-LUX switch functionalized with HSB2.A (head binder) and SD38 (stem binder). B) Titration on H1 from A/Puerto-Rico/8-1934 (H1PR) with a dark-LUX switch functionalized with HSB2.A and SD36. Experimental conditions: Final sensor concentration 300 pM, 1xPBS + 1mg/ml BSA, final NanoGlo dilution of 1:2000, 16 h incubation at 4 °C. Error bars correspond to the standard deviation of n = 3 technical replicates.

**Protein sequences**

The used tags for purifications of proteins are hexa-His-tag (HHHHHH) and StrepTag II (WSHPQFEK).

Dark-LUX A1 scaffold:

Yellow: SmBiT variant, blue: LgBiT, red: DogCatcher; *italic*: L2 linker position; grey: SpyCatcher003; purple: DarkBiT variant.

MGSSHHHHHHSSGLVPRGSHMVTGYRLFEEILGSGGSGGSGGSGGSGGSGSGGSGGSGGSGGSGGSVFTLEDFV
GDWEQTAAYNLDQVLEQGGVSSLLQNLAVSVTPIQRIVRSGENALKIDIHVIIPYEGLSADQMAQIEEVFKVVYPVD
DHHFKVILPYGTLVIDGVTPNMLNYFGRPYEGIAVFDGKKITVTGTLWNGNKIIDERLITPDGSMLFRVTINSSGGGT
KLGEIEFIKVDKTDKKPLRGAVFSLQKQHPDYPDIYGAIDQNGTYQDVRTGEDGKLTFTNLSDGKYRLIENSEPPGYK
PVQNKPIVSFRIVDGEVRDVTSIVPQG*KLGGSGGSGGSGGSGGSGGSGGSGGSGGENLYFQSGGSGGSGGSGGSG*
*GSGGSGGSGGTG*VTTLSGLSGEQGPSGDMTTEEDSATHIKFSKRDEDGRELAGATMELRDSSGKTISTWISDGHVK
DFYLYPGKYTFVETAAPDGYEVATPIEFTVNEDGQVTVDGEATEGDAHTTSGGSSGGSVTGYALFEKESGGSGGSW
SHPQFEK*

A2: SmBiT VTGYRLFEEIL DarkBiT: VTGYALFEKIS

A3: SmBiT: VTGYRLFEKES DarkBiT VTGYALFEKIS

A1 $(TP)_{15}$: L2 position (italic) is GTPTPTPTPTPTPTPTPTPTPTPTPTPTPTPTPG

A1 $(TP)_{22}T$: L2 position (italic) is GTPTPTPTPTPTPTPTPTPTPTPTPTPTPTPTPTPTPTPTPTPTPTG

Dark-LUX 2xDogCatcher A1 scaffold:

Yellow: SmBiT variant, blue: LgBiT, red: DogCatcher; *italic*: L2 linker position; grey: SpyCatcher003; purple: DarkBiT variant.

MGSSHHHHHHSSGLVPRGSHMVTGYRLFEEILGSGGSGGSGGSGGSGGSGSGGSGGSGGSGGSGGSVFTLEDFV
GDWEQTAAYNLDQVLEQGGVSSLLQNLAVSVTPIQRIVRSGENALKIDIHVIIPYEGLSADQMAQIEEVFKVVYPVD
DHHFKVILPYGTLVIDGVTPNMLNYFGRPYEGIAVFDGKKITVTGTLWNGNKIIDERLITPDGSMLFRVTINSSGGGT
KLGEIEFIKVDKTDKKPLRGAVFSLQKQHPDYPDIYGAIDQNGTYQDVRTGEDGKLTFTNLSDGKYRLIENSEPPGYK
PVQNKPIVSFRIVDGEVRDVTSIVPQGKLGGSGGSGGSGGSGGSGGSGGSGGSGGENLYFQSGGSGGSGGSGGS
GGSGGSGGSGGTGKLGEIEFIKVDKTDKKPLRGAVFSLQKQHPDYPDIYGAIDQNGTYQDVRTGEDGKLTFTNLSD
GKYRLIENSEPPGYKPVQNKPIVSFRIVDGEVRDVTSIVPQTSGGSSGGSVTGYALFEKESGGSGGSWSHPQFEK*

HSB2.A-DogTag:

Blue: HSB2.A, green: DogTag.

MGSHHHHHHSSGGIVNVPNCNTTKYQQLARTAVAIYNYHEQAHLTFVENLNCKEQGNYYYITLAATDDAGKKAIY
EAKIGVVESAGWTGVEEFKLVGSGGSGGSGGSGGSGGSSGGSGSGTGDIPATYEFTDGKHYITNEPIPPKGGSGGS
WSHPQFEK*

SD38-DogTag:

Blue: SD38, green: DogTag.

MEVQLVESGGGLVQPGGSLRLSCAVSISIFDIYAMDWYRQAPGKQRDLVATSFRDGSTNYADSVKGRFTISRDNAK
NTLYLQMNSLKPEDTAVYLCHVSLYRDPLGVAGGMGVYWGKGALVTVSSKLGGSGGSGGSGGSGGSTGDIPATY
EFTDGKHYITNEPIPPKGGSGGSWSHPQFEK*

SD38-SpyTag003:

Blue: SD38, green: SpyTag003.

MEVQLVESGGGLVQPGGSLRLSCAVSISIFDIYAMDWYRQAPGKQRDLVATSFRDGSTNYADSVKGRFTISRDNAK
NTLYLQMNSLKPEDTAVYLCHVSLYRDPLGVAGGMGVYWGKGALVTVSSKLGGSGGSGGSGGSGGSTGRGVPHI
VMVDAYKRYKGGSGGSWSHPQFEK

SD36-DogTag:

Blue: SD36, green: DogTag.

MGSVQLVESGGGLVQAGGSLKLSCAASGRTYAMGWFRQAPGKEREFVAHINALGTRTYYSDSVKGRFTISRDNAK
NTEYLEMNNLKPEDTAVYYCTAQGQWRAAPVAVAAEYEFWGQGTQVTVSGGSGGSGTGDIPATYEFTDGKHYIT
NEPIPPKGGSGGSWSHPQFEK*

SD38-slink-DogTag:

MEVQLVESGGGLVQPGGSLRLSCAVSISIFDIYAMDWYRQAPGKQRDLVATSFRDGSTNYADSVKGRFTISRDNAK
NTLYLQMNSLKPEDTAVYLCHVSLYRDPLGVAGGMGVYWGKGALVTVSSKLGGSGGSTGDIPATYEFTDGKHYIT
NEPIPPKGGSGGSWSHPQFEK*

SD38-slink-SpyTag003:

Blue: SD38, green: SpyTag003.

MEVQLVESGGGLVQPGGSLRLSCAVSISIFDIYAMDWYRQAPGKQRDLVATSFRDGSTNYADSVKGRFTISRDNAK
NTLYLQMNSLKPEDTAVYLCHVSLYRDPLGVAGGMGVYWGKGALVTVSSKLGGSGGSTGRGVPHIVMVDAYKRY
KGGSGGSWSHPQFEK*

Library constructs:

Color scheme as in Dark-LUX A1 scaffold, L1,2 and 3 indicating linker positions.

MGGSSHHHHHHSSGLVPRGSHMVTGYRLFEEILGSGGSGGSGGSGGSGGSGSGGSGGSGGSGGSGGSVFTLEDF
VGDWEQTAAYNLDQVLEQGGVSSLLQNLAVSVTPIQRIVRSGENALKIDIHVIIPYEGLSADQMAQIEEVFKVVYPV
DDHHFKVILPYGTLVIDGVTPNMLNYFGRPYEGIAVFDGKKITVTGTLWNGNKIIDERLITPDGSMLFRVTIN-**L1-**
KLGEIEFIKVDKTDKKPLRGAVFSLQKQHPDYPDIYGAIDQNGTYQDVRTGEDGKLTFTNLSDGKYRLIENSEPPGYK
PVQNKPIVSFRIVDGEVRDVTSIVPQ-**L2-**
VTTLSGLSGEQGPSGDMTTEEDSATHIKFSKRDEDGRELAGATMELRDSSGKTISTWISDGHVKDFYLYPGKYTFVE
TAAPDGYEVATPIEFTVNEDGQVTVDGEATEGDAHT-**L3-**VTGYALFEKESGGSGGSWSHPQFEK*
